# The impact of delayed switch to second-line antiretroviral therapy on mortality, depending on failure time definition and CD4 count at failure

**DOI:** 10.1101/661629

**Authors:** Helen Bell-Gorrod, Matthew P Fox, Andrew Boulle, Hans Prozesky, Robin Wood, Frank Tanser, Mary-Ann Davies, Michael Schomaker

**Affiliations:** Health Economics and Decision Science, University of Sheffield, United Kingdom; Center for Global Health and Development, Boston University, Boston, MA, USA; Centre for Infectious Disease Epidemiology and Research, University of Cape Town, Cape Town, South Africa; Division of Infectious Diseases, Department of Medicine, University of Stellenbosch and Tygerberg Hospital, Cape Town, South Africa; Desmond Tutu HIV Centre, Cape Town, South Africa & Institute of Infectious Diseases and Molecular Medicine, University of Cape Town, Cape Town, South Africa; Africa Centre for Health and Population Studies, University of KwaZulu-Natal, Somkhele, South Africa; Institute of Public Health, Medical Decision Making and Health Technology Assessment, UMIT - University for Health Sciences, Medical Informatics and Technology, Hall in Tirol, Austria

**Keywords:** HIV, treatment switching, second-line ART, causal inference, targeted learning

## Abstract

**Background:** Little is known about the functional relationship of delaying second-line treatment initiation for HIV-positive patients and mortality, given a patient’s immune status.

**Methods:** We included 7255 patients starting antiretroviral therapy between 2004-2017, from 9 South African cohorts, with virological failure and complete baseline data. We estimated the impact of switch time on the hazard of death using inverse probability of treatment weighting (IPTW) of marginal structural models. The non-linear relationship between month of switch and the 5-year survival probability, stratified by CD4 count at failure, was estimated with targeted maximum likelihood estimation (TMLE). We adjusted for measured time-varying confounding by CD4 count, viral load and visit frequency.

**Results:** 5-year mortality was estimated as 10.5% (2.2%; 18.8%) for immediate switch and as 26.6% (20.9%; 32.3%) for no switch (49.9% if CD4 count<100 cells/mm^3^). The hazard of death was estimated to be 0.40 (95%CI: 0.33-0.48) times lower if everyone had been switched immediately compared to never. The shorter the delay in switching, the lower the hazard of death, e.g. delaying 30-60 days reduced the hazard 0.52 (0.41-0.65) times, and 60-120 days 0.56 (0.47-0.66) times.

**Conclusions:** Early treatment switch is particularly important for patients with low CD4 counts at failure.

## Introduction

Anti-retroviral treatment (ART) was received by an estimated 4.4 million (61%) people living with HIV in South Africa in 2017^1^. As the number of HIV-positive patients with access to ART has increased, so has the number of patients that have experienced failure of first-line ART. Patients with virological failure on first-line ART should, in principle, switch to second-line therapy as soon as possible, as a delay in switching treatment regimens has been shown to lead to increased mortality ^2–7^. South African guidelines recommend switching from two nucleoside reverse transcriptase inhibitors (NRTIs) and one non-nucleoside reverse transcriptase inhibitor (NNRTI) to two NRTIs and one protease inhibitor (PI) if two consecutive viral loads on first line therapy are greater than 1000 copies/mL. However, in resource limited settings it is still common to delay the switch ^8–10^. Reasons for delays include doubts about adequate patient adherence, availability of viral load testing and the cost of second line regimens^11, 12^.

The effect of delayed switch to second-line therapy on mortality has been investigated in several observational studies which adjusted for measured time-varying confounders using causal inference methods. Gsponer et al. ^5^ showed the drastic reduction in mortality for patients switching to second-line compared to no switch based on an immunological criteria of failing, as well as the benefit of switching early. Petersen et al.^6^ estimated the effect of delayed switch after confirmed virological failure on survival and quantified the relative benefit of earlier switch based on the assumption of a linear relationship between timing of switch and probability of death. Other studies have looked into the impact of delayed switch in South Africa^7^, the effect of using different viral failure definitions^2^ and the relative efficacy of various monitoring strategies^4^.

There have been few studies which have explored the functional relationship between time of switch and mortality^13^, and there is potential for further research into whether there may be a “breaking point” beyond which further delays could be particularly risky, especially for patients with an already compromised immune system. In particular, it would be of interest to know whether the effect of delayed switch is modified by CD4 count at failure. Previous studies have looked at this, albeit in different contexts^6, 7^. Moreover, from a programmatic perspective there may also be a benefit to minimising the time between first viral load greater than 1000 copies/mL and switch given that with new technologies like resistance testing, patients with adequate adherence and proven resistance could potentially be switched earlier. In addition, most of the studies to date had relatively small patient numbers and limited follow-up times.

Our study aims at addressing these gaps. We assess the impact of delayed switch from first-line ART treatment to second-line ART treatment on mortality in 9 South African treatment programs; a large cohort with long follow-up. We use two related but distinct causal approaches; inverse probability of treatment weighting (IPTW) and targeted maximum likelihood estimation (TMLE), which allow us to present or findings on the hazard and incidence scales. The impact of delayed switch is flexibly modelled for patients with different disease severities based on CD4 count at time of viral load failure. We also investigate the importance of monitoring the delay between the first viral load (VL) measure over 1000 copies/ml and confirmed failure (second VL measure >1000 copies /ml) as part of the delay in switch on mortality outcomes.

## Methods

### Study setting and definitions

We included 9 HIV treatment facilities in Southern Africa that took part in the IeDEA-SA collaboration (http://www.iedeasa.org/), namely Desmond Tutu HIV Centre Gugulethu, Hlabisa HIV Treatment and Care Programme, Tygerberg, McCord Hospital, 3 treatment facilities at the Khayelitsha ART Programme, Themba Lethu Clinic and Masiphumelele Clinic. The collaboration has been described in detail elsewhere ^14^.

Adult patients who that started treatment on a first-line treatment regime (2 NRTIs + 1 NNRTI) and failed first-line therapy after 1^st^ January 2004, were included in the analysis. Failure was defined as two consecutive VL measurements greater than 1000 copies/mL and measured at least 4 weeks apart. If measures were taken less than 4 weeks apart the next measure was considered. We excluded patients without any record of receiving ART, those that experienced virological failure within 6 months of ART initiation, those that were not receiving ART at the time of first VL failure and those that switched before viral load failure. In total, we included 7255 patients for the main complete case analysis, see Figure 1, and 8008 patients in the sensitivity analysis with multiple imputation for missing baseline data. Earliest entry date into our sample was 4^th^ October 2004 and the database was closed on 16^th^ August 2017.

**Figure 1:**
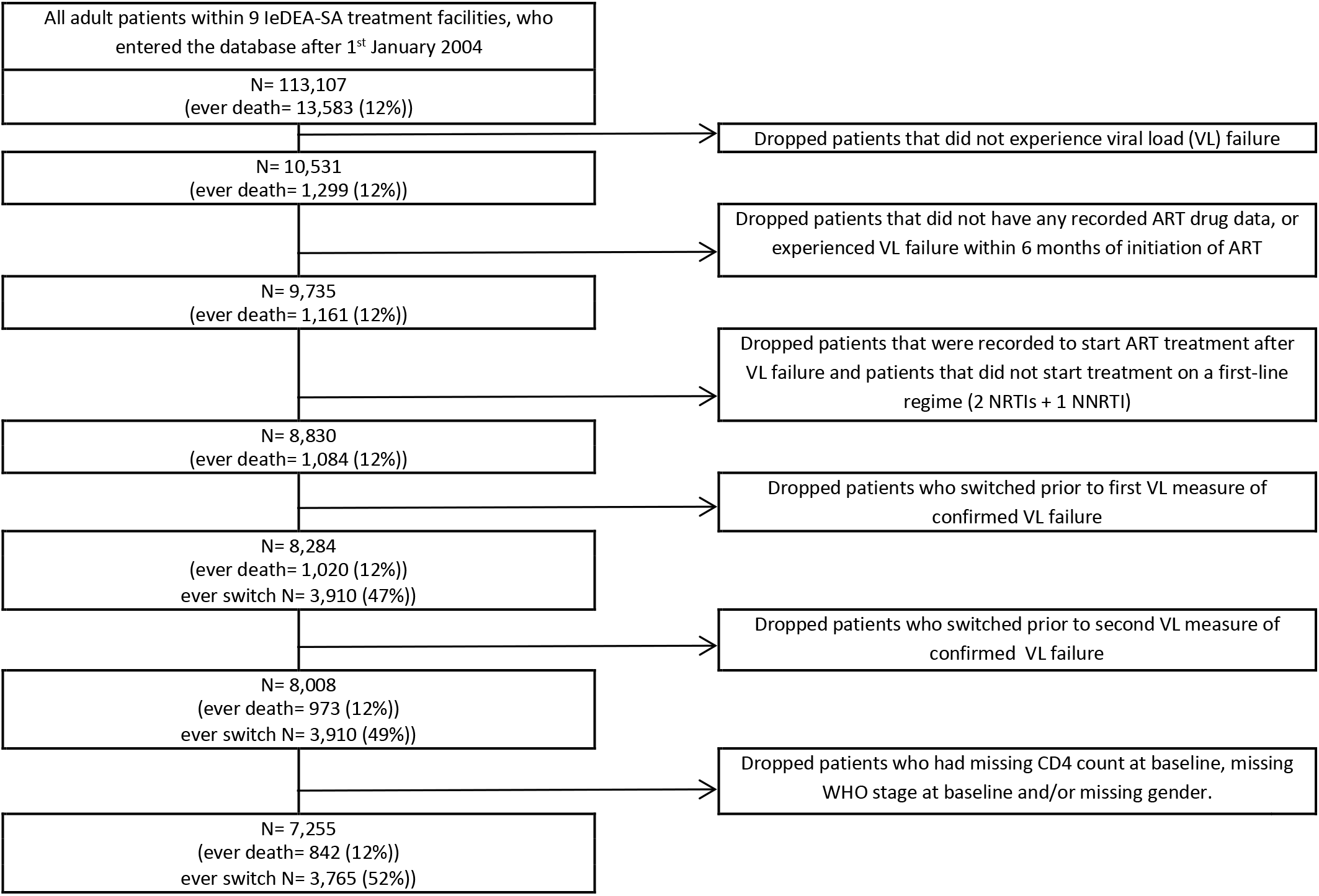
Flow diagram for inclusion of patients in our analysis.

In the main analysis, baseline was defined at the time of first-line viral failure i.e. the date at which the second of the two consecutive viral loads were over 1000 copies per/ml. A secondary analysis was performed using the date at which the first of the two consecutive VLs was greater than 1000 copies per/mL as the baseline, which represents the earliest indication of viral failure. The sample of patients was the same regardless of the definition used because only patients with two elevated viral loads were included. A switch from first-line ART to second-line ART was broadly defined as a switch from 2 NRTIs and 1 NNRTI to 2 NRTIs and 1 PI. A detailed list of second-line regimens in our data is provided in Supplementary Table 1. Patients were defined as being lost to follow-up if there was no visit or event for 9 months after their last recorded visit and before database closure.

The primary endpoint was mortality which was recorded through clinic’s patient files and updated through data from the South African national vital registry where available (this approach is expected to give >96% completeness of mortality data ^15^).

### Analysis

Analysis time started at the date of first-line failure, defined as 2 VL>1000 copies/mL in the main analysis and 1 VL>1000 in the secondary analysis, as described above. Our primary exposure was the timing of switch to second-line ART, measured in months since the respective date of failure and we used this to assess the effect on both the hazard of death and 5-year survival.

Measured and included baseline characteristics (at time of confirmed failure) are age, sex, highest and lowest CD4 count prior to failure, highest and lowest log VL measure prior to failure, an indicator whether a patient was ever suppressed prior to failure, WHO clinical stage at time of ART initiation, year of ART start and treatment facility. Time-varying variables which potentially determined the decision to switch as well as mortality, and were affected by prior treatment regimes, were CD4 count, VL and treatment frequency (measured as number of visits within the past 6 months). It is possible to adjust for confounding of these variables using appropriate causal inference methods ^16^.

We estimated the effect of timing of switch on the hazard of death using inverse probability of treatment weighting (IPTW) of marginal structural models ^2^. To estimate the effect of treatment switch, as well as the non-linear relationship between month since failure and month of switch on the probability of 5-year mortality, stratified by CD4 count at failure, we used targeted maximum likelihood estimation (TMLE) for longitudinal marginal structural working models ^17^.

For IPTW, we used 7 different switching delay strategies; no switch and delayed switch by <30 days, 30-59 days, 60-119 days, 120-179 days, 180-359 days, and ≥ 360 days. We created 7 clones/replicates per patient, one for each treatment strategy, as described previously ^7^. A clone/replicate is censored after it ceases to follow the respective switching strategy. The remaining uncensored observations were weighted to represent what would have happened if the censored patients had continued to follow the respective switching strategy. We used pooled logistic regression models weighted by the stabilized inverse probabilities of treatment and censoring to estimate the effect of the different strategies on the hazard of death. The logistic regression models used to derive the weights contained the above-mentioned time-dependent and baseline variables in the denominator, and baseline variables only in the numerator. The Supplementary Material (Supplementary table 5, Technical Appendix) contains a detailed description of implementation of the method and model specifications. In sensitivity analyses, missing baseline CD4 count and WHO stage were imputed using multiple imputation by chained equations^18^.

With TMLE, we first estimated 5-year mortality under immediate switch after confirmed failure and no switch using the R-package *ltmle* ^19^. The iterated outcome regressions, i.e. the relationship between mortality and the covariates at each point in time (based on 3-month intervals) were estimated using super learning. Super learning is a data-adaptive approach that combines different modelling approaches, such as logistic regression or other regression approaches, such that the expected prediction error (estimated via cross validation) is minimized, see the technical appendix (Supplementary Material) for more details. We then specified marginal structural working models to model the relationship between month since failure, month of switching, and survival, conditional on CD4 count at failure; see technical appendix and the footnote in Figure 3 for more details. The fitted models, calculated based on the approach described in Petersen et al. ^17^, were then used to visualize the relationship.

All analyses were conducted in Stata 13 ^20^ and R 3.5.1 ^21^.

### Ethics

This IeDEA-SA collaboration study was approved by the University of Cape Town and University of Bern human research ethics committees. At most sites, the requirement for informed consent was waived, as only anonymized data that were already collected as part of routine monitoring contributed to the collaborative dataset.

## Results

Median time from ART start to failure was 1218 days (about 3.3 years); median time from confirmed failure to switch was 121 days (1^st^ quartile: 49 days; 3^rd^ quartile: 288 days), with follow-up times from confirmed failure ranging between 1 and 4409 days (median 1835 days, IQR 1183-2470). During follow-up 3765 patients (52%) switched, and 842 (12%) died.

The included patients were mostly female (65%), and had advanced WHO stage at ART initiation (60%), see Table 1. Among patients that never switched, a substantial proportion (19%) had a viral load >100.000 copies/mL at confirmed viral load failure.

**Table 1:**
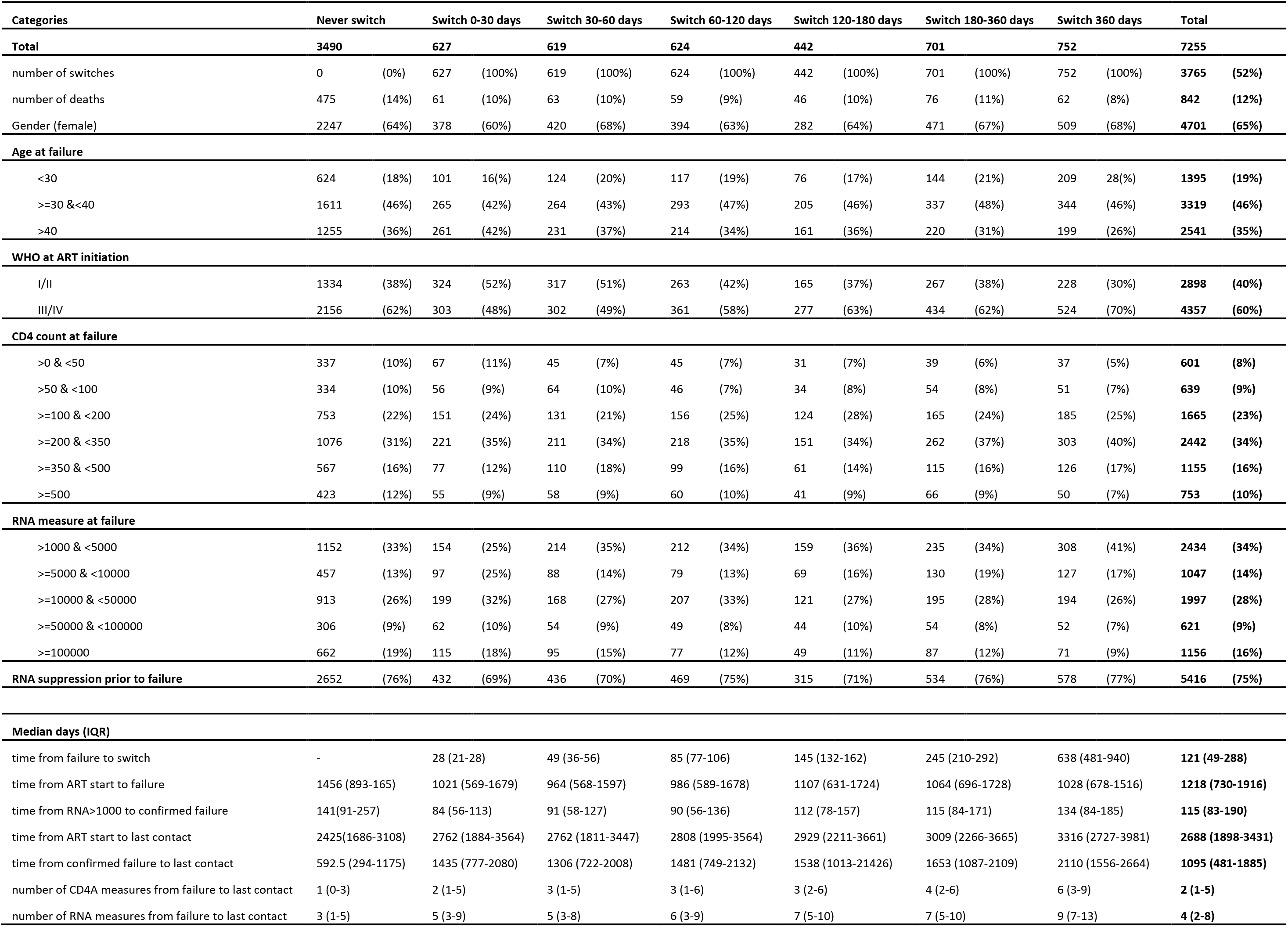
Characteristics of patients at confirmed viral load failure (second consecutive viral load measure greater than 1000 copies/ml)

The probability of being switched was higher among patients with low current CD4 count, high VL, and a higher visit frequency (Table 2). These variables also predicted the probability of death, confirming that they are likely time-varying confounders.

**Table 2:**
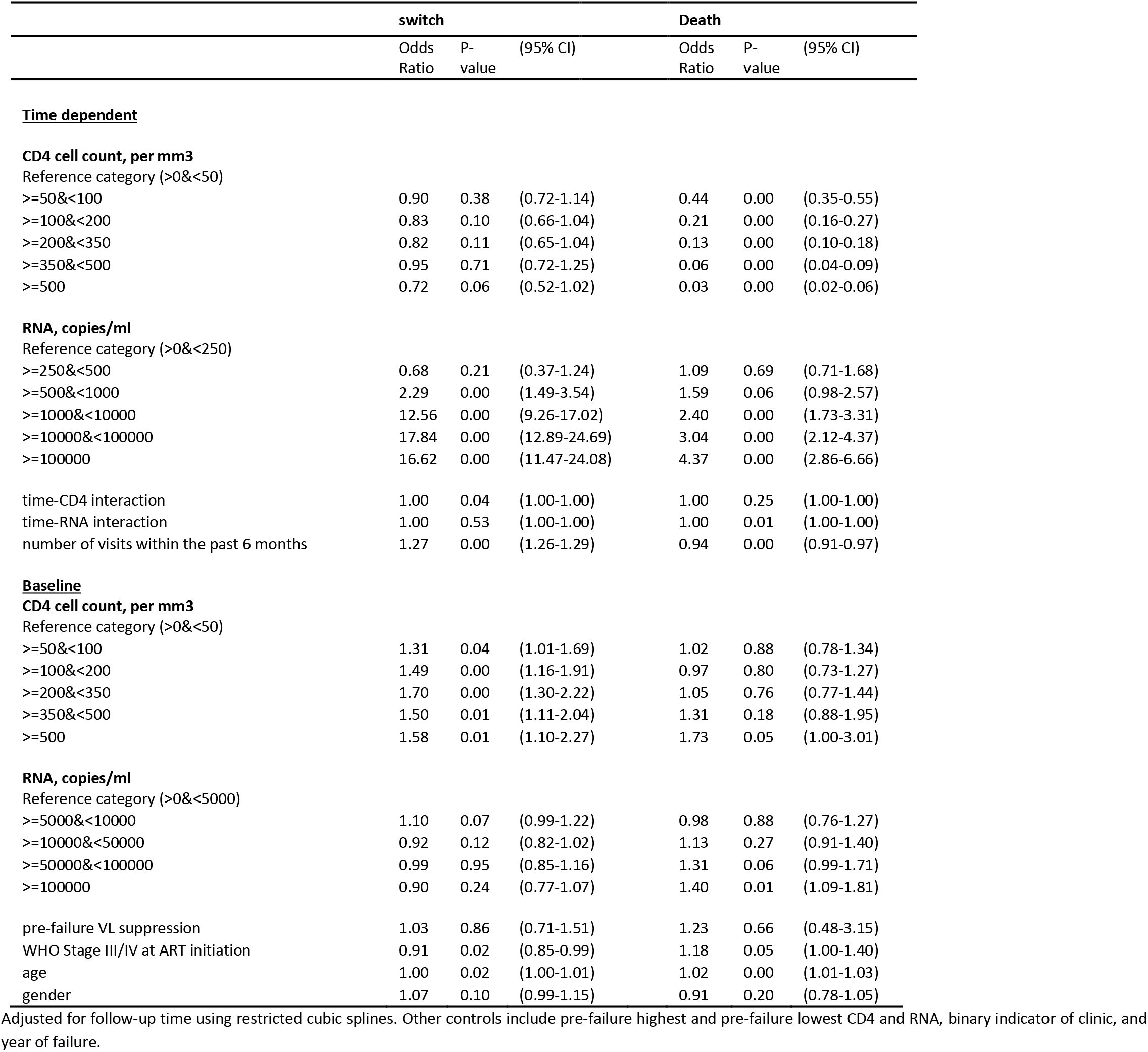
Predictors of switch from first-line to second-line ART and predictors of death.

The effect of immediate switch compared to no switch on mortality, if confirmed failure was used as failure definition, was estimated as 0.49 (95% CI: 0.42-0.58) in a crude analysis, and as 0.37 (0.30-0.46) using IPTW. Results with multiple imputation were 0.47 (0.40-0.54) in a crude analysis, and 0.36 (0.30-0.44) using IPTW. If first VL>1000 copies/mL was used as definition of failure the estimates were 0.52 (0.45-0.61) and 0.42 (0.34-0.52) respectively. After imputation the results were 0.50 (0.43-0.58) and 0.41 (0.34-0.51) (Supplementary Table 2). Figure 2 shows that the shorter the delay in switching, the lower the hazard of death. There are stronger benefits of early switch when considering one VL>1000 copies/mL as failure definition. Similar results are obtained after multiple imputation of baseline CD4 count and WHO stage (Supplementary Table 2). Sensitivity analyses show that truncation of the stabilized weights at the 1^st^ and 99^th^ quantile yields the most stable results (Supplementary Table 3).

**Figure 2:**
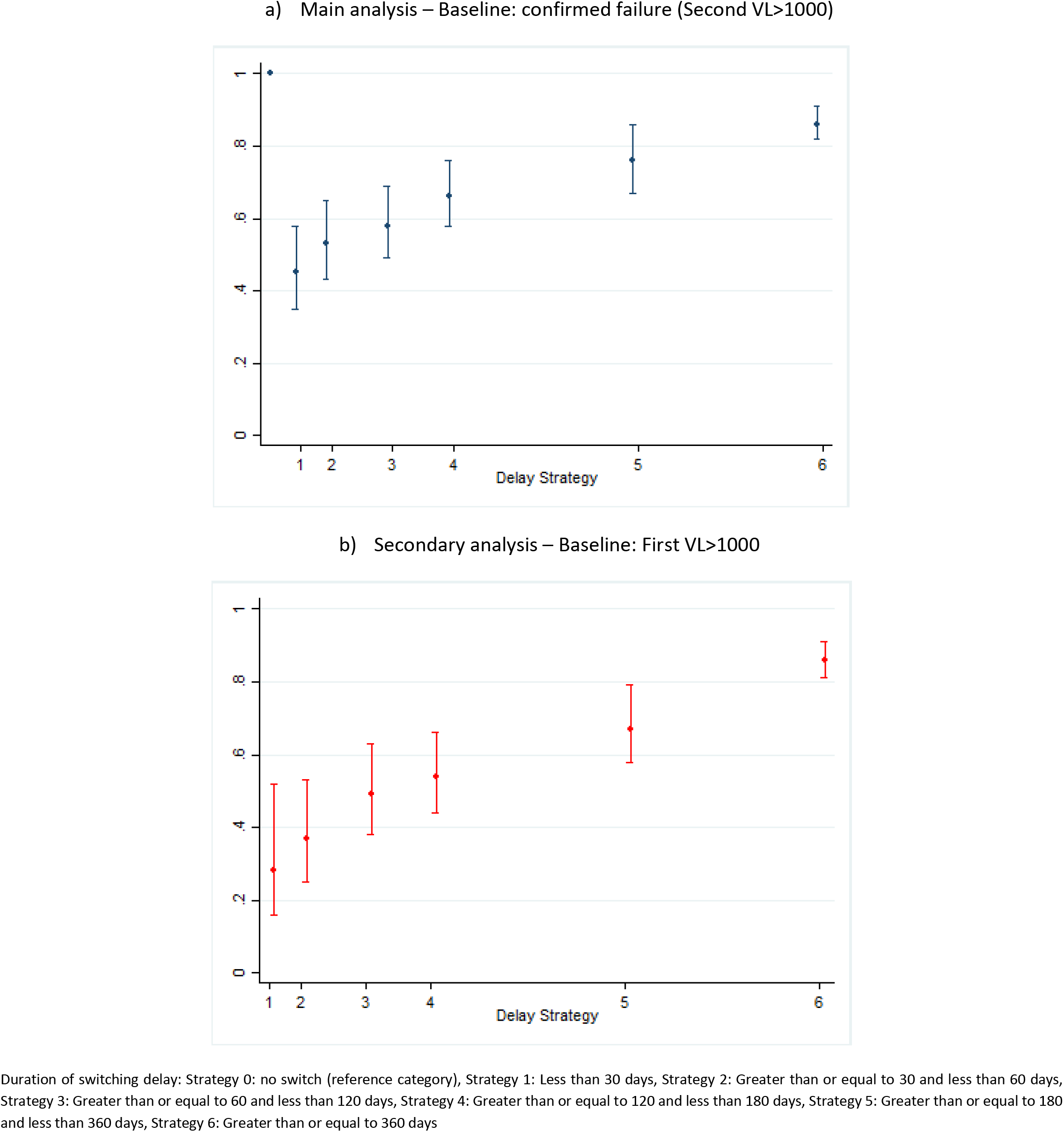
Hazard ratio of each switching delay duration subgroup vs no switch using IPW of MSM.

Using TMLE, 5-year mortality was estimated as 10.5% (2.2%; 18.8%) if everyone had been switched immediately, and as 26.6% (20.9%; 32.3%) if everyone had stayed on their failing regimen. The corresponding risk difference was −16.1% (−26.1%; −6.1%), and the odds ratio was 0.32 (0.13; 0.82). The working MSM’s, fitted with TMLE, are visualized in Figure 3. The black dashed line shows that the estimated 5-year mortality (i.e. 60 months after failure) to be about 25% under no switching (month of switch = 60). However, this varies considerably by immune status at failure. Almost 51% would have died among those who had a CD4 count <100 at failure (red line), but only a small proportion (17.5%) among those with a CD4 count > 200 cells/mm^3^ (green line). Moreover, the effect of delaying treatment was more severe (i.e. steeper ascent) among patients failing with CD4 count < 100 cells/mm^3^. Similar results are obtained when evaluating probabilities of death <5 years (Supplementary Figure 1). Overall, the estimated relationship between switch time and mortality was non-linear, as visualized in Figure 3. This is because the estimated coefficients of the non-linear switch time terms in the working MSMs were important, and also significant at the 5% level.

**Figure 3:**
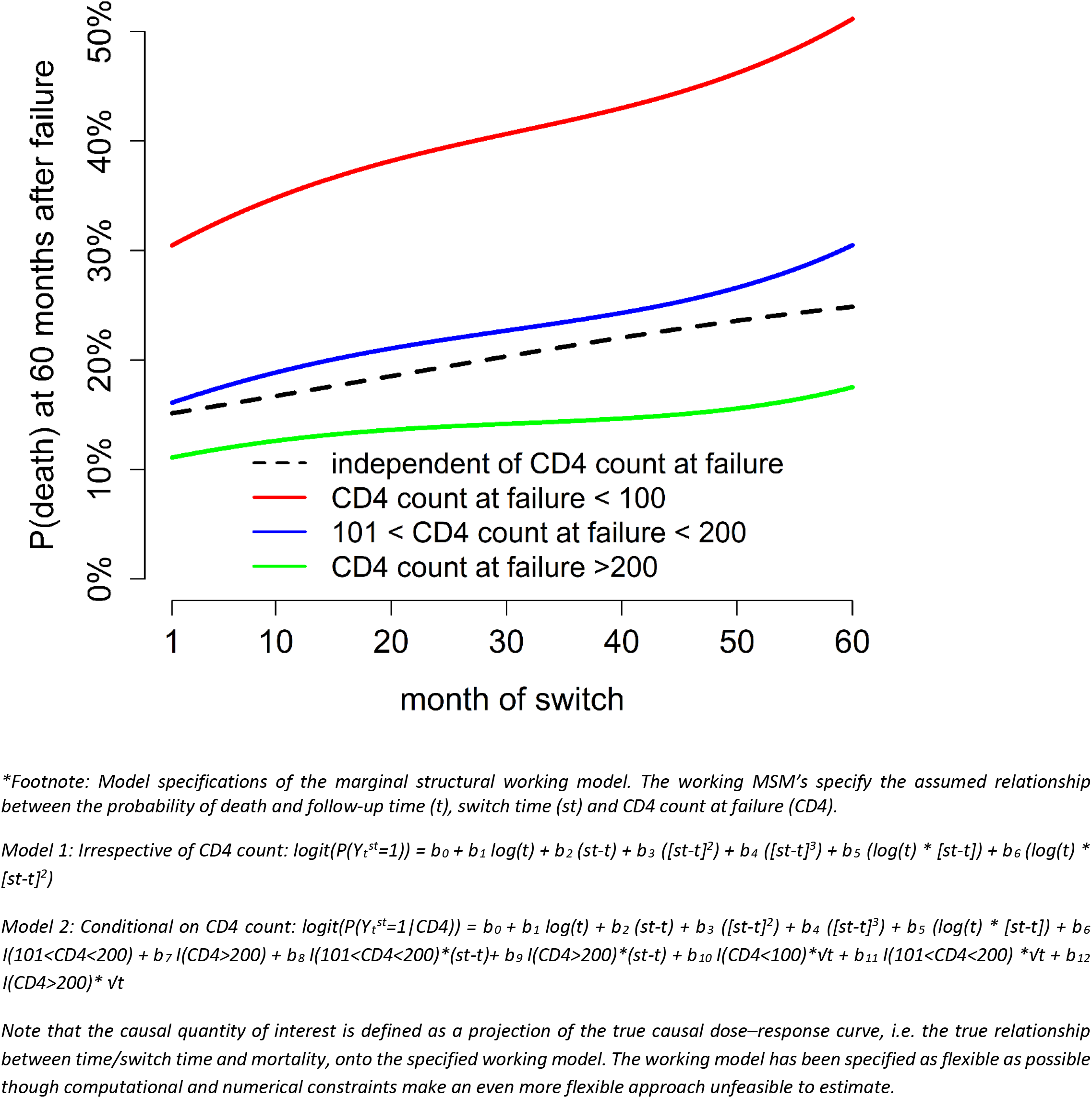
Probability of death 5 years after virologic failure, for different CD4 count categories at time of failure, and depending on month of switch (i.e. extent of delay). Estimates are based ‘on working MSM’s estimated with LTMLE as specified under the footnote*.

## Discussion

### Statement of principal findings

Our study highlights that it often takes a long time to switch patients to second line treatment in Southern Africa. We have shown that an early switch of regimen is highly beneficial in terms of reduced mortality. Patients with low CD4 counts at time of failure are at particularly high risk of increased mortality, whereas a moderate delay in healthy patients comes with a comparatively lower risk.

### Strengths and limitations

Our study is based on a large data set, with a multitude of different treatment regimens and long follow-up, which allowed us to model the relationships in the data in a flexible and robust way. Since our patients have relatively regular viral load measurements for the setting, we have been able to evaluate the effect of switching based on viral failure, rather than immunological failure; which is of great interest given that viral load monitoring is typically not available in public sector programs in resource limited settings, though it is currently being expanded. Another strength is the use of causal inference methods to adjust for time-dependent confounding affected by prior treatment, which would not be possible with traditional regression analyses ^16^. This helped us to contrast switching strategies under different viral failure definitions. We also used TMLE, which has desirable statistical properties (double robustness), to confirm and extend the MSM analysis. In contrast to previous studies, we have even been able to implement this method for a marginal structural model that postulated non-linear relationships between treatment strategies and survival.

Our study has some limitations. Our analysis is based on routine data from South African treatment programs. It may well be possible that patients defined to be lost to follow-up are in fact cycling in and out of care, possibly in different provinces ^22^; or that the complication of capturing start and stop dates of different drugs may lead to inaccuracies that could potentially also affect our ability to accurately define switch dates. The diagnostics further suggested that there could be some positivity violations in our data which means that individuals may not have a positive probability of continuing to receive treatment according to a specific treatment rule, given that they have done so thus far and irrespective of the covariate history (Supplementary Table 4, Supplementary Figure 2). This could have affected our estimates. Another limitation is the unavailability of patient-level adherence data.

There are additional limitations associated with the first VL>1000 at baseline (secondary) analysis, which occur due to the definition of the sample. Eligibility for the sample is based on confirmed failure. After first VL>1000, those included cannot switch or die until after their next VL measurement, thus creating a period of immortal time. Table 1 indicates that the period of time between first VL>1000 and confirmed failure is greater, on average, for those with longer delays between confirmed failure and switch. Hence, this may cause some bias in the comparisons of delay strategies. Furthermore, the restriction of the first VL>1000 sample to patients that attained confirmed failure (VL>1000) at next VL measurement means that the secondary analysis can only be interpreted in reference to the confirmed-failure population, and therefore is not generalizable to the wider population.

### Interpretation of findings

It is no surprise that delayed treatment switch may affect patient’s health. However, according to our results, earlier switch is of particular benefit when switching after the first sign of failure, i.e. the first viral load > 1000 copies/mL, for those that go on to confirmed failure. HIV specialists may be reluctant to switch patients that have adherence problems or are unstable, but for stable patients who fail because of resistance or toxicities, early switching after a first elevated viral load could be of benefit.

Our results confirm that switching is partly determined by visit frequency, which may relate to clinician concern for patients based on health status, but also strongly relates to patient’s engagement in care and adherence. To reduce the risk of failure of another regimen, patients on second-line treatment should be adherent. We have shown the benefit of switching even under imperfect adherence, but ideally patients should be psychologically prepared to adhere to their new treatment regimen.

### Results in context

Our results comparing immediate switch to no switch yield similar conclusions to other studies which used other definitions of failure, which were done in different patient populations, for different follow-up times, and used different methodological approaches^5–7, 17^. Like Rohr et al.^7^ we show the that the effectiveness of switching strategies depends on disease severity, though in a more refined way given that we modelled the relationship non-linearly for different patient groups. Similar to other studies we have shown that remaining on first-line therapy leads to an increase in mortality compared to switching, and that earlier switch is beneficial in terms of survival ^6, 17^. Our marginal structural working models were more complex than the MSMs in these studies, which makes a more refined interpretation of the dose-response relationship between delay in switching and mortality possible; however, both previous studies^13^ and current research^23^ suggests that it may be important to allow for even more flexible approaches to model specification and fitting than ours. Nevertheless, whatever methodological approach is chosen, it is important to note that the beneficial effect of switching can be observed for different definitions of treatment failure ^5, 6^.

Our results have two direct implications for current programme guidance. Firstly, for stable virologically suppressed patients, it is no longer recommended in South Africa that they receive regular CD4 counts. However, once a patient is viraemic, our results demonstrate the critical importance of CD4 count in further risk stratifying patients. The value of dropping routine CD4 count testing in the interests of cost-saving, needs to be considered alongside the benefits of the additional information it provides on disease severity and mortality risk, and could be used to highlight groups that are in more urgent need of early switch.

In patients who subsequently fail virologically, we have demonstrated that the delay between the first and second elevated viral load contribute to the non-linear early increase in mortality resulting from delayed switching, especially in patients with low CD4 counts. This points to the importance of either accelerating confirmation of virological failure in patients with advanced immunological suppression, or to consider switching at the first evidence of viraemia if cost and regimen-sparing are no longer important considerations driving the need to confirm virologic failure.

### Further research

In the South-African context, and according to WHO guidelines, switching is permitted after confirmed failure. Hence, our analyses were restricted to a subgroup of patients with 2 consecutive VL>1000. The wider dataset, indicated in figure 1, shows that some patients switch onto second-line treatment prior to confirmed virologic failure. It would be interesting to investigate the impact of time to switch from first elevated VL using a sample defined with the eligibility criteria of one VL>1000. In this larger sample, the additional complication of the competing risk of virologic re-suppression would need to be considered in the analysis, as re-suppressing patients would no-longer be eligible for switch.

## Conclusions

Our study highlights the importance of early treatment switch, particularly for patients with low CD4 counts at failure.

## Acknowledgements

This project was supported by Grant Number U01AI069924 from NIH (NIAID, NICHD, NCI, NIDA, NIMH), (PI: Egger and Davies). Its contents are solely the responsibility of the authors and do not necessarily represent the official views of the NIH. We thank Maya Petersen for feedback on an earlier version of our LTMLE analysis and Julia Rohr for feedback on our IPTW analysis.

## Supplementary Material: Technical appendix

*Notation:* Let Y_t_ be the binary survival outcome measured at time t, 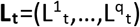 the vector of time-varying covariates at time t (CD4 count, log_10_ viral load, visit frequency [number of visits within the past 6 months]), C_t_ a censoring indicator at time t, and A_t_ (antiretroviral) treatment at time t. The follow-up time is t=0,1,3,6,9,…,60 months. **L_0_** is the vector of baseline covariates which contains are age, sex, highest and lowest CD4 count prior to failure, highest and lowest log VL measure prior to failure, an indicator whether a patient was ever suppressed prior to failure, WHO clinical stage, year of ART start and treatment facility, CD4 count, visit frequency and viral load. We are interested in the intervention vector A=(a_0_,a_1_,a_3_,a_6_,…,a_60_) which is a multiple-time point intervention where at each time point antiretroviral therapy may be given or not. More generally, we refer to the intervention history (up to and including time t) as 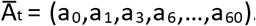. For example, immediate treatment initiation refers to 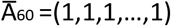 and no treatment initiation to 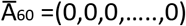. With the superscript we denote counterfactuals. For example, 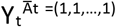 is the outcome that would have been observed at time t had everyone received (possibly contrary to the fact) immediate treatment initiation, i.e. ART at all time points. A rule *d* assigns treatment A_t_ such that it starts at a specific time point (and therefore determines the amount of delay in treatment initiation). Since the rule effectively determines the switch time (*st*) we write Y^st^ to refer to the outcome that would have been observed under a rule that assigns treatment in line with a certain switch time.

### Target Quantities

We are interested in estimating

i. how the counterfactual probability of death 60 months after first-line failure varies as a function of the assigned switch time (*st*); where switch time based on rule *d* determines how the treatment vector A looks like. That is, we are interested in
  a. 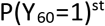 where switch time *st* varies between 0 and 60 months; and
  b. a marginal structural working model to summarize how the counterfactual probability of death at follow-up time *t* varies as a function of *t* and assigned switch time (*st*) [and therefore treatment vector A]; see below for the model specification.
ii. We are also interested in summarizing the effect of the delay strategy *d* on (the hazard λ of) mortality with marginal structural Cox models of the form

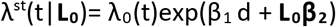 The Cox model is approximated with a pooled logistic regression model containing (splines of) follow-up time and the (above mentioned) baseline variables **L_0_**. The exact model specification is given further below.

### Structural assumptions

As Petersen et al.^1^ we assume that CD4 count, HIV RNA (viral load) and clinic visit frequency influence decisions whether and when to switch therapy, as well as affecting mortality; since these variables are affected by prior switching decisions and mediate the effect of exposure to failing first-line therapy on mortality, *standard regression adjustment methods* are not suitable, see below for our estimation approaches. We speculate that (unmeasured) patient adherence also affects decisions of when to switch, as well as mortality.

### Observed data & Identification

Our data contains O=(**L**,A,Y,C) as defined above under “notation”. The target quantities above [listed under i) and ii)] can be identified under the assumptions of sequential randomization (“no unmeasured confounders”), consistency (“well-defined intervention”) and positivity^2, 3^. With positivity we mean that a patient who has not already switched should have some positive probability of both switching and not switching (regardless of his covariate history). With no unmeasured confounding, we essentially mean that those variables that affect the decision of when to switch and mortality, and are themselves affected by prior treatment decisions, are all contained in **L_t_** (see above point on adherence). More formal definitions of the above assumptions are given in Petersen et al. ^4^ and Schomaker et al. ^5^.

### Estimation

To estimate the target quantities listed in i) we use longitudinal targeted maximum likelihood estimation as described in Petersen et al. ^4^ and implemented in the R-package *ltmle* and for ii) we use inverse probability weighting of marginal structural models following the approach in Rohr et al.^6^, see also Cain et al.^7^ for more details.

<u>For estimation of the targeted quantities in i)</u> we follow exactly the approach described in detail in Petersen et al. ^4^. Briefly, we estimate P(Y_t_=1|**L_0_**)^st^ for all possible switch times (i.e. delay strategies *d* that delay treatment by *st* = 0,1,3,6,9,…,60 months) and follow-up times *t*=0,1,3,6,9,…,60 and summarize the dose-response relationship between Y and *t* and *st* in two different working models:

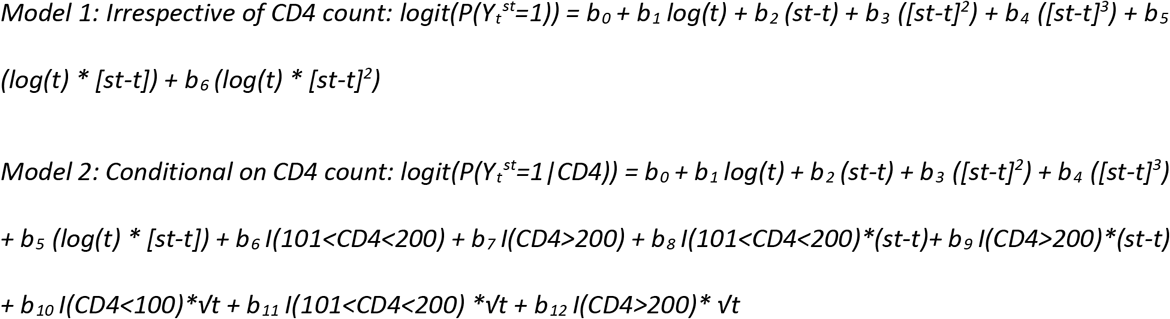

Model 1 summarizes the dose-response relationship independent of CD4 count at time of viral failure; model 2 summarizes the relationship conditional on CD4 count (at time of failure). The transformations for follow-up time have been chosen such that the working MSM yields similar results as the (non-MSM) estimates for the probability of death at 5 years under 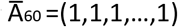 and 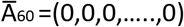 respectively. The working models allow for an inflection point in the survival curve with respect to time of switch due to the inclusion of a cubic polynomial of difference in switch time and follow-up time. Unfortunately, more complex working models that include switch time in a way that is more sophisticated could not be fitted due to technical constraints. Figure 3 and supplementary Figure 1 have been produced based on the estimates of the above working MSMs.

To estimate the above target quantity i)a), we have used the ltmle() function in the package *ltmle*. With this we estimated that 5-year mortality was 10.5% (2.2%; 18.8%) if everyone had been switched immediately, and as 26.6% (20.9%; 32.3%) if everyone had stayed on their failing regimen. The corresponding risk difference was −16.1% (−26.1%; −6.1%), and the odds ratio was 0.32 (0.13; 0.82).

To estimate the target quantity i)b), we used the function ltmleMSM() in *ltmle*. To estimate the conditional (nested) outcome expectations needed for both a) and b), as well as the treatment and censoring models/mechansism we used super learning as recommended previously^8^. In more detail, we used the following learners: the arithmetic mean, (generalized linear) regression models with all main terms [GLM], GLMs based on an EM-algorithm-Bayesian model fitting, GLMs chosen by stepwise variable selection with Akaikes Information Criterion [AIC], GLMs containing interaction terms, as well as additive models; these learners have been partly fitted on the whole set of covariates as well as subsets based on screening with Cramer’s V ^9^ and Lasso estimation ^10^.

<u>For estimation of the targeted quantities in ii),</u> we estimated the effect of immediate switch compared to no switch on mortality, if confirmed failure was used as failure definition, as 0.37 (0.30-0.46) using IPTW. If first VL>1000 copies/mL was used as definition of failure the estimates were 0.42 (0.34-0.52) respectively. For both analyses we needed models for the treatment and censoring mechanisms to calculate weights, for each patient at each time period. We applied stabilised weights (as defined and explained below) which require estimation of a numerator and a denominator. The models we need are as follows:

### Models for the treatment (and artificial censoring in the delay analysis) mechanism

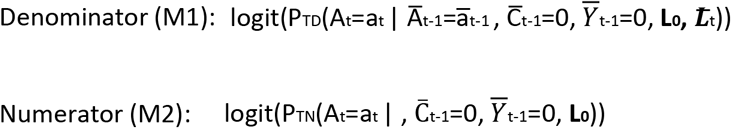

#### Model specification

Baseline covariates **L_0_** included in the model were binary indicators of baseline CD4 (>=50 & <100, >=100 & <200, >=200 & <350, >=350 & <500, >=500) and binary indicators of baseline viral load (>250 & <500, >=500 & <1000, >=1000 & <10000, >=10000 & <100000, >=100000) as well as age, gender, clinic, and binary indicators for calendar year of failure (2003-2006, 2007-2009, 2010-2012, 2013-2017). We also included a binary indicator of pre-failure VL suppression and categorical variables for pre-failure highest and pre-failure lowest CD4 and RNA. Time dependent variables **L_t_** included binary indicators of categorical CD4 and viral load, linear CD4-time and viral load-time interactions, and number of visits within the past 6 months. Supplementary Table 5 lists the fitted models in detail.

### Models for the loss-to-follow censoring mechanism

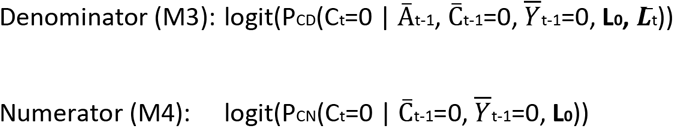

#### Model specification

The model specifications for the censoring mechanisms included **L_0_** and **L_t_** as described above in the treatment models, except that we excluded the (linear) time-CD4 and time-RNA interactions. Supplementary Table 5 lists the fitted models in detail.

### Stabilised weights

<u>For the simple “switch versus no switch” analysis</u> (presented in manuscript text and in supplementary tables 2 and 3), treatment and censoring stabilised weights were derived from denominator and numerator of the treatment and loss-to-follow-up censoring models M1-M4. These weights were combined, as follows, to create a combined treatment and censoring stabilised weight, for each person, at each time point.

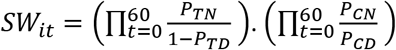

<u>For the “delay in switch versus no switch” analysis</u> (presented in Figure 2), we followed the approach in Rohr et al.^6^ (and Cain et al.^7^) and described in the manuscript. Again, models for the treatment and censoring mechanisms were required and fitted in line with the model specifications given above. Treatment and censoring weights were estimated prior to the expansion and artificial censoring of the dataset.

The dataset was expanded by replicating each person-time observation 6 times to create a set of 7 clones; one clone to represent each of the 7 delay regimes. Within each regime, person time was artificially censored according to adherence to the delay regime. For instance, a person that switched at 75 days from baseline would be censored at 30 days in the switch within 30 days regime, would be censored at 60 days in the 30-59 days regime, would not be censored in the 60-119 days regime, and would be censored at 75 days in the 120-179 days, 180-359 days, greater than 360 days and never switch regimes. The cloning allowed one person to follow multiple regimes simultaneously, therefore estimates become more efficient,^7^.

For the treatment weights, probabilities from the treatment models were used to generate weights for each person-time-regime observation. First, probabilities were assigned to each observation based on the following rules, where P_T_ represents the probabilities derived from the numerator/denominator treatment model;

- Treatment rules; P_T_ at time of switch, 1 after switch, and 1-P_T_ before switch
- Artificial censoring rules; 1-P_T_ if at time of artificial censoring if it is also time of switch, and P_T_ at time of artificial censoring if it is not also time of switch.

Second, cumulative probabilities for numerator and denominator were calculated for each person over time within each regime. Third, stabilised treatment weights were estimated using the cumulative numerator and denominator probabilities for each person at time point, for each regime. This follows the approach described in the supplementary materials of Cain, et al^26^.

For loss-to-follow-up censoring, stabilised censoring weights were created using cumulative probabilities from the numerator and denominator lost-to-follow-up censoring models. Treatment and loss-to-follow-up stabilised weights were combined to create a combined treatment and censoring stabilised weight, for each person, at each time point, within each regime.

Stabilized weight summaries are given in Supplementary Table 3.

- Marginal structural Cox models were fitted in line with the model specification given in ii) under the above heading “Target quantities”, based on weighted pooled logistic regression.
  For the “switch versus no switch analysis”;
  Marginal Structural Cox model: logit(P(Y_t_=1| A_t_, **L_0_**))
  For the “delay in switch versus no switch analysis”;
  Marginal Structural Cox model:
  logit(P(Y_t_=1 | d_t_, **L_0_**))

d*_t_* indicates a set of binary variables which represent the delay strategies/regimes. Baseline covariates L_0_ included in the two models above were binary indicators of baseline CD4 (>=50 & <100, >=100 & <200, >=200 & <350, >=350 & <500, >=500) and binary indicators of baseline viral load (>250 & <500, >=500 & <1000, >=1000 & <10000, >=10000 & <100000, >=100000) as well as age, gender, clinic, and binary indicators for calendar year of failure (2003-2006, 2007-2009, 2010-2012, 2013-2017). We also included a binary indicator of pre-failure VL suppression and categorical variables for pre-failure highest and pre-failure lowest CD4 and RNA. Confidence intervals were calculated using cluster robust standard error estimators.

### Diagnostics

The diagnostics for IPTW of marginal structural models are summarized in supplementary Table 3. For the LTMLE analyses we provide the percentage of truncated cumulative inverse treatment and censoring probabilities. We used a truncation level of 1%. Large percentages of truncation suggest limited data support for these interventions and possible positivity violations^5^. The working MSM is meant to extrapolate well for interventions where there is little data support^4^. The summary (rounded percentages) is as follows:

**Supplementary Table 4:** LTMLE truncation.

| Delay (months) | 0 | 1 | 3 | 6 | 9 | 12 | 15 | 18 | 21 | 24 | 27 |
| --- | --- | --- | --- | --- | --- | --- | --- | --- | --- | --- | --- |
| % truncation | 2 | 0 | 0 | 1 | 4 | 12 | 11 | 8 | 11 | 23 | 14 |
| Delay (months) | 30 | 33 | 36 | 39 | 42 | 45 | 48 | 51 | 54 | 57 | 60 |
| % truncation | 27 | 22 | 28 | 13 | 16 | 25 | 29 | 18 | 25 | 75 | 1 |

One can see the limited data support for intervention strategies that delay switching by 12-57 month.

Note however that these interventions have a lower impact on the fitted working MSM ^16^. It is however important to stress that even under an MSM approach estimates remain vulnerable to positivity violations^11^. A particular concern is that standard errors may be anti-conservative, though recent developments suggest that it is possible to construct estimators that are somewhat more robust with respect to positivity violations.^12^

The distributions of the fitted cumulative inverse treatment and censoring probabilities after 5 years of follow-up are visualized in Supplementary Figure 2. One can again see the limited data support for intervention strategies that delay switching by 12-57 month.

## Supplementary Material

**Supplementary Table 1:** Drug regimens for second-line treatment.

| Regime at switch | Frequency | Percent | Cum. |
| --- | --- | --- | --- |
| 3TC KLT TDF | 1,232 | 31.51 | 99.92 |
| 3TC AZT KLT | 932 | 23.84 | 57.11 |
| AZT DDI KLT | 760 | 19.44 | 20.2 |
| 3TC AZT KLT TDF | 125 | 3.20 | 61.05 |
| AZT KLT RTV | 92 | 2.35 | 22.92 |
| 3TC D4T KLT | 83 | 2.12 | 65.52 |
| 3TC D4T EFV KLT TDF | 77 | 1.97 | 63.38 |
| 3TC AZT EFV KLT TDF | 74 | 1.89 | 33.22 |
| KLT RTV TDF | 67 | 1.71 | 27.11 |
| 3TC ABC KLT | 51 | 1.3 | 28.67 |
| FTC KLT TDF | 36 | 0.92 | 24.78 |
| 3TC D4T KLT NVP TDF | 29 | 0.74 | 66.29 |
| 3TC AZT KLT NVP TDF | 24 | 0.61 | 57.85 |
| 3TC KLT SQV | 22 | 0.56 | 68.41 |
| 3TC AZT D4T EFV KLT | 21 | 0.54 | 30.36 |
| KLT RTV SQV | 18 | 0.46 | 25.4 |
| AZT KLT TDF | 17 | 0.43 | 23.35 |
| 3TC D4T KLT TDF | 15 | 0.38 | 66.68 |
| 3TC EFV KLT TDF | 14 | 0.36 | 67.26 |
| 3TC ATV AZT | 11 | 0.28 | 28.98 |
| ABC KLT RTV | 10 | 0.26 | 0.33 |
| 3TC ATV TDF | 10 | 0.26 | 29.64 |
| 3TC D4T EFV KLT NVP TDF | 10 | 0.26 | 61.41 |
| 3TC KLT RTV | 10 | 0.26 | 67.85 |
| Other* | 170 | 4.00 | 100 |
| Total | 3,910 | 100 |  |

\*Includes combinations of drugs that include 3TC, ATV NVP, D4T, DDI, EFV, TDF, ABC, FTC, RTV, DRV, RGV, KLT, ETV, NFV, SQV

| Type of treatment | Abbreviation | Treatment name |
| --- | --- | --- |
| NRTI | 3TC | Lamivudine |
| NRTI | ABC | Abacavir |
| NRTI | AZT | Zidovudine |
| NRTI | D4T | Stavudine |
| NRTI | DDC | Zalcitabine |
| NRTI | DDI | Didanosine |
| NRTI | FTC | Emtricitabine |
| NRTI | TDF | Tenofovir |
| NRTI/NNRTI | ATP | Atripla |
| NNRTI | EFV | Efavirenz |
| NNRTI | ETV | Etravirine |
| NNRTI | NVP | Nevirapine |
| NNRTI | RPV | Rilpivirine |
| PI | ATV/r | Atazanavir/Ritonavir |
| PI | DRV | Darunavir) |
| PI | LPV/r | Lopinavir/Ritonavir |
| PI | NFV | Nelfinavir |
| PI | RTV | Ritonavir |
| PI | SQV | Saquinavir |
| PI | TPR | Tipranavir |
| CCR 5 antagonist | MVC | Maraviroc |
| InSTI | RGV | Raltegravir |

**Supplementary Table 2:** Complete case analysis and results after multiple imputation of missing baseline data of WHO stage at ART initiation and CD4 count.

| Baseline | Main analysis<br>Second VL measurement<br>of Confirmed failure |  | Secondary analysis<br>First VL measurement<br>of Confirmed failure |  |
| --- | --- | --- | --- | --- |
|  | HR | 95% CIs | HR | 95% CIs |
| **Time from baseline to switch\*** |  |  |  |  |
| **Results from complete case analysis** |  |  |  |  |
| Crude (switch vs no switch) | 0.49 | (0.42-0.58) | 0.52 | (0.45-0.61) |
| IPTW (switch vs no switch) | 0.37 | (0.30-0.46) | 0.42 | (0.34-0.52) |
| <u>IPW (delay in switch vs no switch):</u> |  |  |  |  |
| No switch (reference category) | 1 | - | 1 | - |
| Less than 30 days | 0.45 | (0.35-0.58) | 0.28 | (0.16-0.52) |
| Greater than or equal to 30 and less than 60 days | 0.53 | (0.43-0.65) | 0.37 | (0.25-0.53) |
| Greater than or equal to 60 and less than 120 days | 0.58 | (0.49-0.69) | 0.49 | (0.38-0.63) |
| Greater than or equal to 120 and less than 180 days | 0.66 | (0.58-0.76) | 0.54 | (0.44-0.66) |
| Greater than or equal to 180 and less than 360 days | 0.76 | (0.67-0.86) | 0.67 | (0.58-0.79) |
| Greater than or equal to 360 days | 0.86 | (0.82-0.91) | 0.86 | (0.81-0.91) |
| **Results after multiple imputation of missing variables** |  |  |  |  |
| Crude (switch vs no switch) | 0.47 | (0.40-0.54) | 0.50 | (0.43-0.58) |
| IPTW (switch vs no switch) | 0.36 | (0.30-0.44) | 0.41 | (0.34-0.51) |
| <u>IPW (delay in switch vs no switch):</u> |  |  |  |  |
| No switch (reference category) | 1 | - | 1 | - |
| Less than 30 days | 0.43 | (0.34-0.55) | 0.33 | (0.18-0.62) |
| Greater than or equal to 30 and less than 60 days | 0.51 | (0.42-0.61) | 0.39 | (0.26-0.58) |
| Greater than or equal to 60 and less than 120 days | 0.58 | (0.50-0.68) | 0.50 | (0.39-0.63) |
| Greater than or equal to 120 and less than 180 days | 0.64 | (0.56-0.73) | 0.51 | (0.42-0.62) |
| Greater than or equal to 180 and less than 360 days | 0.72 | (0.64-0.81) | 0.65 | (0.56-0.76) |
| Greater than or equal to 360 days | 0.88 | (0.83-0.92) | 0.87 | (0.83-0.91) |

\*Note that in the first VL measurement analysis, an additional 30 days from baseline to the upper and lower limits of each delay category were included to account for the fact that patients in our sample were not permitted to switch until 4 weeks after first VL measure >1000copies/ml. Crude refers to a switch vs no switch analysis without inverse probability weighting.

**Supplementary Table 3:** stabilized weights diagnostics for the switch versus no switch analysis.

| Truncation percentiles | Estimated weights for the outcome Death |  | Estimates of effect of switch on Death |  |
| --- | --- | --- | --- | --- |
|  | Mean (SD) | Minimum/maximum | Hazard/Odds ratio | Standard Error |
| **Baseline: Confirmed failure (Main analysis)** |  |  |  |  |
| 100 | 1.07 (1.82) | 0.04/119.75 | 0.31 | 0.05 |
| 99.5 | 1.02 (0.99) | 0.04/10.71 | 0.37 | 0.04 |
| 99 | 0.98 (0.68) | 0.04/5.36 | 0.37 | 0.04 |
| 97.5 | 0.94 (0.47) | 0.04/2.67 | 0.38 | 0.04 |
| 95 | 0.91 (0.38) | 0.04/1.82 | 0.38 | 0.04 |
| 90 | 0.88 (0.32) | 0.04/1.43 | 0.39 | 0.04 |
| **Baseline: First VL>1000 copies/mL (Secondary analysis)** |  |  |  |  |
| 100 | 5692.25 (3940592) | 0.02/3.247e+09 | + | + |
| 99.5 | 1.00 (1.03) | 0.02/11.01 | 0.42 | 0.05 |
| 99 | 0.97 (0.78) | 0.02/6.75 | 0.42 | 0.05 |
| 97.5 | 0.91 (0.44) | 0.02/2.57 | 0.42 | 0.04 |
| 95 | 0.88 (0.36) | 0.02/1.75 | 0.42 | 0.04 |
| 90 | 0.85 (0.31) | 0.02/1.35 | 0.42 | 0.04 |

+ indicates that the outcome model did not converge.

**Supplementary Table 5:** Output from the treatment and censoring models.

|  | Treatment – 2 VL>1000 |  |  |  | Censor – 2 VL>1000 |  |  |  | Treatment – VL>1000 |  |  |  | Censor – VL>1000 |  |  |  |
| --- | --- | --- | --- | --- | --- | --- | --- | --- | --- | --- | --- | --- | --- | --- | --- | --- |
|  | denominator |  | numerator |  | denominator |  | numerator |  | denominator |  | numerator |  | denominator |  | numerator |  |
|  | Odds Ratio | P-value | Odds Ratio | P-value | Odds Ratio | P-value | Odds Ratio | P-value | Odds Ratio | P-value | Odds Ratio | P-value | Odds Ratio | P-value | Odds Ratio | P-value |
| **Time dependent** |  |  |  |  |  |  |  |  |  |  |  |  |  |  |  |  |
| **CD4 cell count, per mm3** |  |  |  |  |  |  |  |  |  |  |  |  |  |  |  |  |
| >=50<<100 | 0.90 | 0.38 | - | - | 1.23 | 0.01 | - | - | 1.07 | 0.52 | - | - | 1.17 | 0.03 | - | - |
| >=100<<200 | 0.83* | 0.10 | - | - | 1.25 | 0.00 | - | - | 0.94 | 0.54 | - | - | 1.21 | 0.00 | - | - |
| >=200<<350 | 0.82 | 0.11 | - | - | 1.37 | 0.00 | - | - | 0.98 | 0.85 | - | - | 1.33 | 0.00 | - | - |
| >=350<<500 | 0.95 | 0.71 | - | - | 1.37 | 0.00 | - | - | 1.09 | 0.46 | - | - | 1.33 | 0.00 | - | - |
| >=500 | 0.72* | 0.06 | - | - | 1.43 | 0.00 | - | - | 0.81 | 0.14 | - | - | 1.33 | 0.00 | - | - |
| **RNA, copies/ml** |  |  |  |  |  |  |  |  |  |  |  |  |  |  |  |  |
| >250<<500 | 0.68 | 0.21 | - | - | 0.92 | 0.08 | - | - | 0.72 | 0.28 | - | - | 0.93 | 0.17 | - | - |
| >=500<<1000 | 2.29*** | 0.00 | - | - | 0.82 | 0.00 | - | - | 2.30 | 0.00 | - | - | 0.85 | 0.01 | - | - |
| >=1000<<10000 | 12.56*** | 0.00 | - | - | 0.71 | 0.00 | - | - | 12.01 | 0.00 | - | - | 0.78 | 0.00 | - | - |
| >=10000<<100000 | 17.84*** | 0.00 | - | - | 0.67 | 0.00 | - | - | 16.67 | 0.00 | - | - | 0.74 | 0.00 | - | - |
| >=100000 | 16.62*** | 0.00 | - | - | 0.69 | 0.00 | - | - | 15.18 | 0.00 | - | - | 0.78 | 0.00 | - | - |
| time-CD4 interaction | 1.00** | 0.04 |  |  |  |  |  |  | 1.00 | 0.04 |  |  |  |  |  |  |
| time-RNA interaction | 1.00 | 0.53 | - | - | - | - | - | - | 1.00 | 0.92 | - | - | - | - | - | - |
| number of visits within the past 6 months | 1.27*** | 0.00 | - | - | - | - | - | - | 1.33 | 0.00 | - | - | - | - | - | - |
| **Baseline** |  |  |  |  |  |  |  |  |  |  |  |  |  |  |  |  |
| **CD4 cell count, per mm3** |  |  |  |  |  |  |  |  |  |  |  |  |  |  |  |  |
| >=50<<100 | 1.31** | 0.04 | 1.09 | 0.38 | 1.01 | 0.85 | 1.08 | 0.30 | 0.95 | 0.66 | 0.89 | 0.21 | 0.99 | 0.88 | 1.03 | 0.74 |
| >=100<<200 | 1.49*** | 0.00 | 1.32 | 0.00 | 0.98 | 0.82 | 1.08 | 0.25 | 1.14 | 0.21 | 1.02 | 0.81 | 0.91 | 0.16 | 0.97 | 0.69 |
| >=200<<350 | 1.70*** | 0.00 | 1.51 | 0.00 | 1.04 | 0.61 | 1.19 | 0.01 | 1.17 | 0.17 | 1.08 | 0.39 | 0.94 | 0.41 | 1.05 | 0.48 |
| >=350<<500 | 1.50*** | 0.01 | 1.42 | 0.00 | 1.03 | 0.73 | 1.20 | 0.01 | 0.99 | 0.95 | 0.94 | 0.55 | 0.90 | 0.16 | 1.01 | 0.94 |
| >=500 | 1.58*** | 0.01 | 1.21 | 0.09 | 0.90 | 0.21 | 1.07 | 0.41 | 1.08 | 0.62 | 0.80 | 0.05 | 0.78 | 0.00 | 0.88 | 0.11 |
| **RNA, copies/ml** |  |  |  |  |  |  |  |  |  |  |  |  |  |  |  |  |
| >=5000<<10000 | 1.10* | 0.07 | 1.21 | 0.00 | 0.99 | 0.74 | 0.97 | 0.47 | 0.90 | 0.04 | 1.03 | 0.59 | 1.05 | 0.19 | 1.04 | 0.34 |
| >=10000<<50000 | 0.92 | 0.12 | 1.26 | 0.00 | 1.06 | 0.07 | 1.03 | 0.34 | 0.89 | 0.02 | 1.08 | 0.07 | 1.02 | 0.66 | 1.00 | 0.97 |
| >=50000<<100000 | 0.99 | 0.95 | 1.35 | 0.00 | 1.03 | 0.53 | 1.00 | 0.93 | 0.78 | 0.00 | 0.97 | 0.70 | 1.00 | 0.98 | 0.98 | 0.65 |
| >=100000 | 0.90 | 0.24 | 1.12 | 0.07 | 0.99 | 0.87 | 0.97 | 0.51 | 0.64 | 0.00 | 0.77 | 0.00 | 1.04 | 0.34 | 1.02 | 0.65 |
| pre-failure VL suppression | 1.03 | 0.86 | 1.01 | 0.96 | 1.06 | 0.69 | 1.02 | 0.88 | 1.21 | 0.33 | 1.06 | 0.77 | 1.00 | 0.98 | 1.00 | 0.97 |
| WHO Stage III/IV | 0.91** | 0.02 | 0.96 | 0.29 | 1.06 | 0.03 | 1.05 | 0.12 | 0.90 | 0.00 | 0.94 | 0.12 | 1.06 | 0.05 | 1.04 | 0.16 |
| age | 1.00** | 0.02 | 1.00 | 0.15 | 1.00 | 0.56 | 1.00 | 0.99 | 1.00 | 0.03 | 1.00 | 0.08 | 1.00 | 0.82 | 1.00 | 0.40 |
| gender | 1.07* | 0.10 | 1.07 | 0.07 | 1.04 | 0.19 | 1.06 | 0.04 | 1.08 | 0.03 | 1.09 | 0.03 | 1.07 | 0.03 | 1.08 | 0.01 |

All models were adjusted for follow-up time using restricted cubic splines, and included binary categorical variables for pre-failure highest and pre-failure lowest CD4 and RNA, binary indicator of clinic, and binary indicator of year of failure (2003-2006, 2007-2009, 2010-2012, 2013-2017). VL suppression was defined as RNA below 400 copies per ml

**Supplementary Figure 1:**
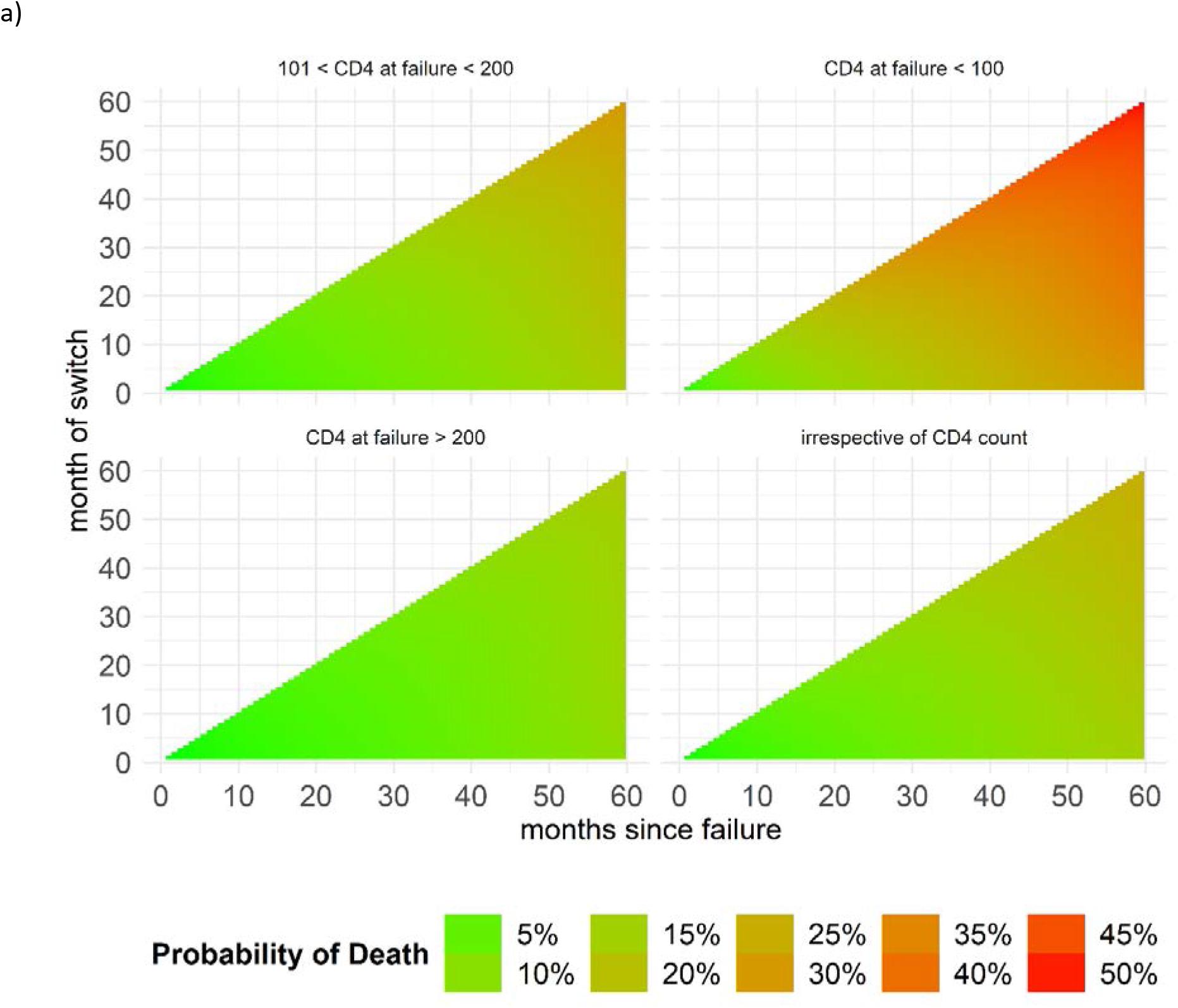
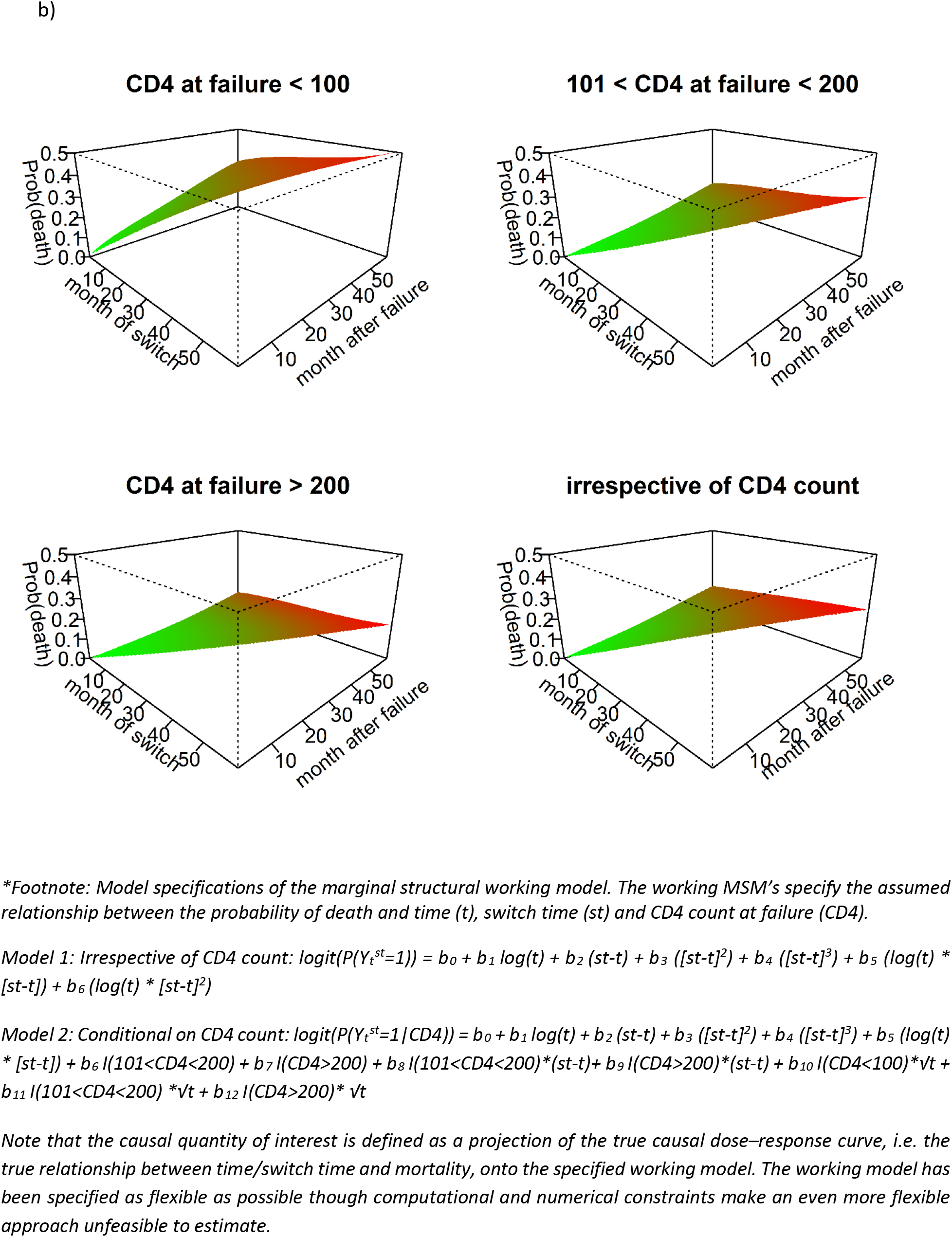
Probability of death 5 years after virologic failure, for different CD4 count categories at time of failure, and depending on month after failure and month of switch. Estimates are based on a working MSM as specified in the technical appendix and under the footnote*. Panel a) visualizes the results in a contour plot where the probability of death is represented by colours and panel b) plots the probability of death in a third dimension, on the z-axis. Note that for both a) and b) the curves at 60 months after failure equate to the results plotted in Figure 3 in the main text. Red colours refer to higher probabilities of death.

**Supplementary Figure 2:**
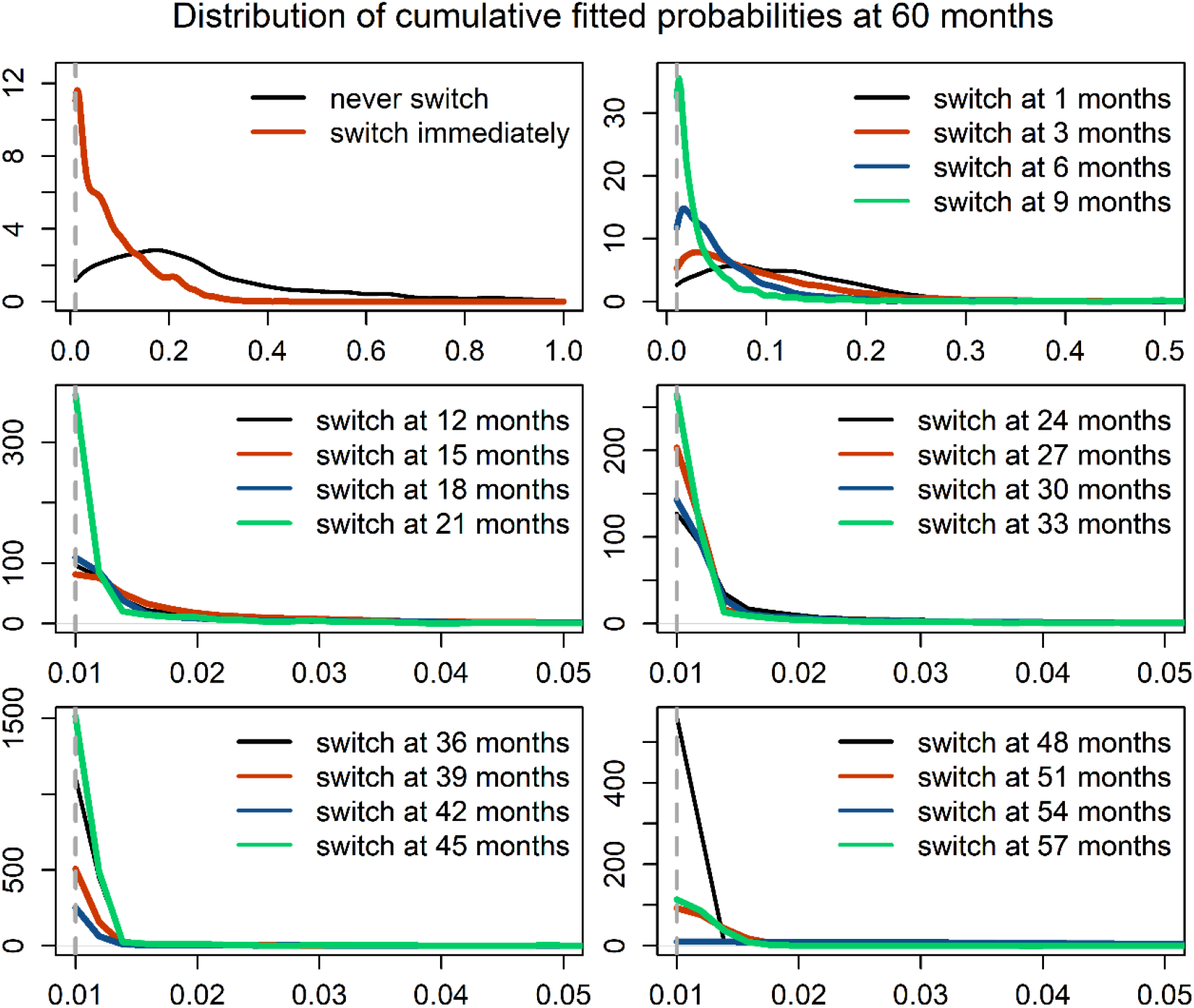
Kernel density plots of the distribution of cumulative fitted probabilities (after 60 months of follow-up) for the different switch strategies.

